# Inverted trophic pyramids and transient fish community reorganization in large reservoirs

**DOI:** 10.64898/2026.09.21.753197

**Authors:** Alejandro Sepúlveda-Correa, Marco A. Rodríguez, Katrine Turgeon

## Abstract

The rapid global expansion of dams has significantly reshaped freshwater ecosystems, yet long-term trophic dynamics and food-web stability within reservoirs remain underexplored. This study examines the reorganization of fish communities induced by the trophic surge following dam construction and river impoundment. We used the predator-to-consumer mass ratio (PCMR) as a proxy for trophic pyramid structure to track changes in fish biomass distribution across three large boreal reservoirs in Quebec, Canada, over 20 years, using data from before and after impoundment. We found transient top-heaviness events (*i.e.*, PCMR values > 1; predator accumulation), characterized by a hump-shaped trajectory of the PCMR values in two reservoirs (Opinaca and Robert Bourassa). The third reservoir (Caniapiscau) showed a gradual increase in top-heaviness but did not exhibit a clear transient pattern. The magnitude and duration of top-heaviness varied across reservoirs, potentially due to differences in the system’s capacity to receive and accumulate nutrients, which are driven by water residence time and reservoir size. This research provides a conceptual framework for understanding the long-term trophic dynamics of reservoirs, which can help inform the development of more effective management strategies to preserve ecological health in these ecosystems.

## INTRODUCTION

The growing demand for water and energy is driving rapid infrastructure development globally, with dams increasingly altering rivers and streams by obstructing their natural flows to create artificial reservoirs (Gielen et al. 2019). Globally, there are now at least 50,000 large dams (over 15 meters in height) and nearly a million smaller impoundments, which collectively modify the flow of about two-thirds of the water that would otherwise reach the oceans (Nilsson et al., 2005; Lehner et al., 2024). Extensive research on large-scale transformations of the biotic and abiotic components of freshwater and riparian ecosystems has highlighted the profound impacts of damming (Baxter, 1977; Friedl & Wüest, 2002; Nilsson et al., 2005; Poff et al., 2007; Turgeon et al., 2019a; Monaghan et al., 2020). Most studies agree that the primary negative impact of dams is the disruption of natural hydrological regimes. However, the trophic responses triggered by impoundment and dam operations remain underexplored due to limited data, increasing uncertainty about their enduring effects on aquatic ecosystems. As reservoir development continues, understanding these ecological dynamics is essential for creating management strategies that sustain ecosystem health and biodiversity.

During the first years following impoundment, reservoirs are dynamic environments in which the interplay between bottom-up and top-down regulation unfolds in often predictable ways. The trophic surge hypothesis (TSH) provides a framework for understanding the initial ecological dynamics of reservoirs (Baranov, 1961). According to the TSH, reservoirs experience a substantial nutrient pulse from the flooded watershed, resulting in a primary productivity surge (Ostrofsky & Duthie, 1980; Turgeon et al., 2016). This influx of nutrients acts as a “biological windfall”, sparking cascading effects throughout the food web and eventually reaching the highest trophic levels (Turgeon et al., 2016; Bumpers et al., 2017; Monaghan et al., 2020).

As time progresses, reservoirs undergo a phase of trophic depression in which the initial nutrient pulse is depleted, resulting in a distinct hump-shaped trajectory in primary production over time (Paterson et al., 2019; Trottier et al., 2024). As bottom-up regulation weakens, top-down effects take precedence, shifting the direction of energy flow. The once-thriving higher trophic levels, which had flourished in the nutrient-rich environment, face food scarcity, setting off cascading effects throughout the food web (McQueen et al., 1989). Although an equilibrium state is expected to emerge eventually, this stabilization can be delayed by dam operations (*e.g.*, drawdown and cyclical filling), which may hinder the establishment of a steady state and prolong trophic rearrangements across food web compartments (Turgeon et al., 2016). The boost of trophic dynamics in reservoir ecosystems is driven by an initial disturbance, the nutrient “pulse,” which is followed by periodic, smaller-scale “press” disturbances that lead to constant, smaller trophic rearrangements (Bender et al., 1984). These surge–depression dynamics can alter ecological traits and processes that have evolutionarily shaped species’ adaptations and trophic interactions (Poff et al., 2007). Consequently, species within reservoirs are expected to modify their interactions and distributions across both spatial and temporal scales (Grill et al., 2015), leading to major food-web rearrangements, with new structures and dynamics emerging at every trophic level (Cross et al., 2013; Mor et al., 2018).

Energy transfer efficiency is conventionally assumed to be approximately 10%, which should result in bottom-heavy Eltonian pyramids (Elton, 1927; Lindeman, 1942). Inverted trophic pyramids (ITPs) challenge this model because higher-level biomass exceeds that of lower levels (Sandin & Zgliczynski, 2015; McCauley et al., 2018). Although some ITPs may stem from methodological biases such as predator overestimation (Ward-Paige et al., 2010; Trebilco et al., 2013), compelling evidence confirms that ITPs can occur under specific natural conditions (Trebilco et al., 2016; McCauley et al., 2018). Beyond their unusual trophic structure, ITPs can amplify cascading effects and increase susceptibility to instability if prey availability fails to meet predator demand (Rip & McCann, 2011; McCauley et al., 2018). Nevertheless, top-heavy communities may persist and remain stable when sustained by high basal turnover rates, energy subsidies from spatial boundaries, or maintained energy flow efficiency (Del Giorgio & Gasol, 1995; Gellner et al., 2016; Heathcote et al., 2016; Vadeboncoeur & Power, 2017).

Significant shifts in fish distribution and community composition in reservoirs are driven by a complex interplay of factors, including food availability, competition, predator-prey imbalances, and species-specific functional traits (Brosse et al., 2007; Agostinho et al., 2008, 2016; Prchalová et al., 2009; Říha et al., 2009; Turgeon et al., 2019a, 2019b; Perônico et al., 2020). These responses vary widely based on limnological and geophysical characteristics of the reservoir such as area, depth, volume, and water residence time, as well as individual species’ adaptations to new conditions (Fee, 1979; Cohen & Newman, 1988; Post et al., 2000; Håkanson, 2005). Consequently, some fish communities exhibit resilience or transient imbalances linked to the TSH, while others show little evidence of long-term declines in productivity (Gido et al., 2000; Perônico et al., 2020; Pereira et al., 2021; Parisek et al., 2024).

The analysis of ITPs goes beyond simply describing biomass typologies within communities and seeks to describe the energetic dynamics and mechanisms that enhance energy transfer efficiency, shaping species interactions in reservoirs. We focus here on the following question: if trophic surges lead to improved productivity at higher trophic levels, how will the fish community respond once the initial nutrient pulse dissipates? Specifically, we consider whether the trophic surge following impoundment unbalances the top predator-to-prey ratio and leads to the establishment of ITPs in reservoir fish communities. We analyze the conditions that could lead to a true inversion of the trophic pyramid, or to top-heavy configurations which do not necessarily represent a complete inversion in biomass distribution.

Our research objectives were to: 1) assess the plausibility of TSH as a key mechanism driving top-heavy configuration in reservoirs by using temporal time series of the predator-to-consumer mass ratio (PCMR) to track post-impoundment changes in fish biomass distribution, and 2) review the main mechanisms influencing the trophic balance of fish communities following dam construction, with a particular emphasis on water residence time (WRT) and reservoir size. We also present a conceptual framework, grounded in our empirical results, to better understand the long-term ecological dynamics of large reservoirs.

## METHODS

### Study sites

The La Grande Rivière hydroelectric complex (LG complex) is a system of cascade reservoirs built between 1979-1996 to harness the hydroelectric potential of major rivers in northern Quebec, Canada. The creation of the LG complex resulted in the formation of seven large reservoirs by diverting three major rivers: the Caniapiscau (flow reduced by 43% at its mouth), the Eastmain (86% reduction), and the Opinaca (86% reduction; Therrien et al., 2002). Fish communities were systematically monitored by Hydro-Québec in three key reservoirs before and after impoundment: Robert-Bourassa (RB; impounded in 1979), Opinaca (OP; 1980), and Caniapiscau (CA; 1982) (Table 1; Fig. 1).

**Figure 1.**
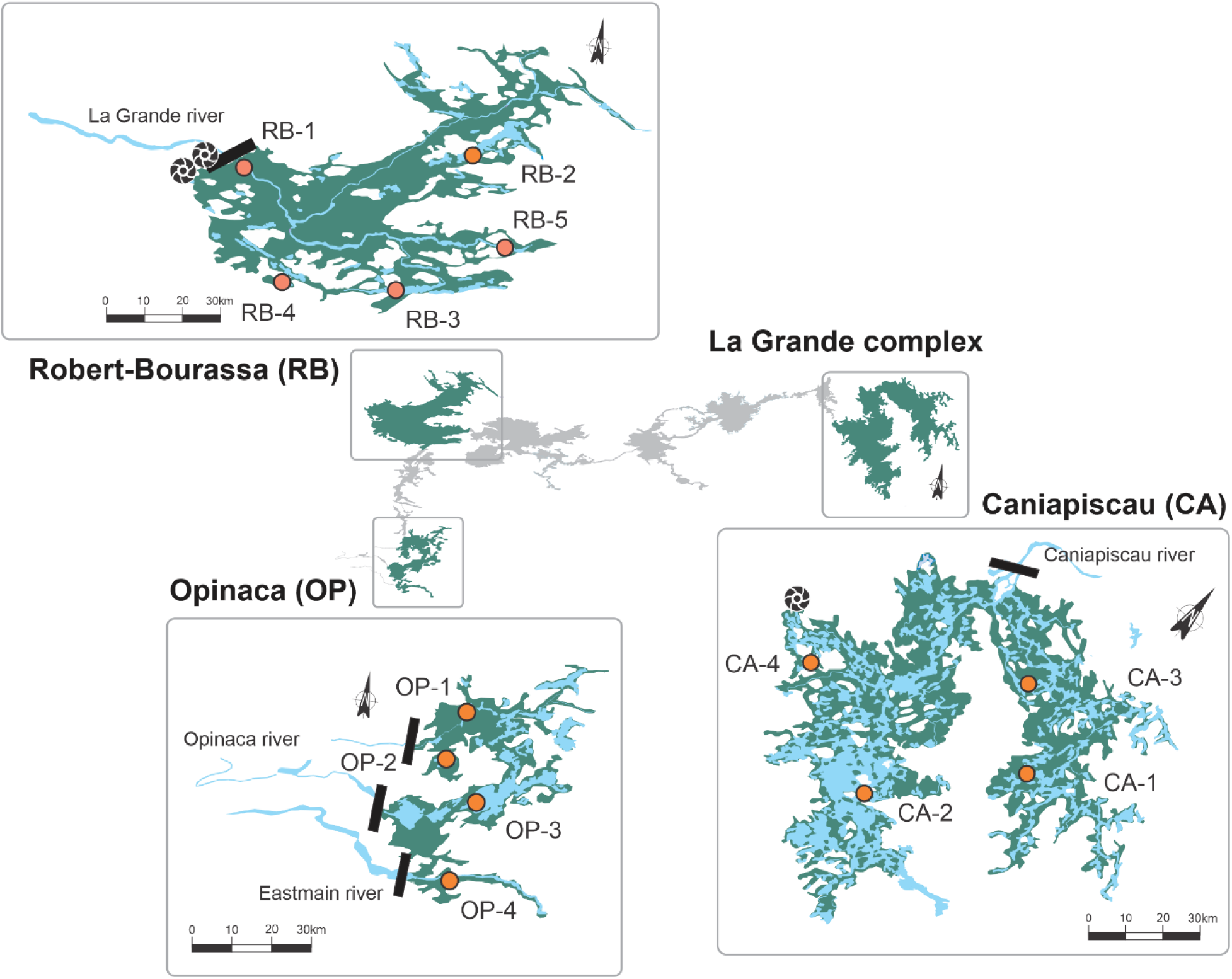
Maps of the three reservoirs included in this study: Robert-Bourassa, Opinaca, and Caniapiscau, located within the La Grande hydroelectric complex in northern Quebec. Sampling stations (orange points) are identified as RB-1 to RB-5 in Robert-Bourassa, OP-1 to OP-4 in Opinaca, and CA-1 to CA-4 in Caniapiscau. The extent of the waterbodies is shown prior to impoundment (light blue) and after impoundment (green).

**Table 1.** Geographical, physical, and limnological characteristics of the Robert-Bourassa, Opinaca, and Caniapiscau reservoirs in northern Quebec. All three reservoirs are oligotrophic and were impounded within three years. The reservoirs differ notably in size (area, volume, depth) and hydrological dynamics (water residence time and mean annual drawdown).

| Variable | Reservoirs |  |  |
| --- | --- | --- | --- |
|  | Opinaca | Robert-Bourassa | Caniapiscau |
| Latitude | 52°38'58''N | 53°45'00''N | 54°31'46''N |
| Longitude | 76°19'54''W | 77°00'00''W | 69°51'18''W |
| Trophic status | Oligotrophic | Oligotrophic | Oligotrophic |
| Area of the reservoir (km<sup>2</sup>) | 1040 | 2835 | 4275 |
| Area flooded (km<sup>2</sup>) | 740 | 2630 | 3430 |
| Volume of the reservoir (km<sup>3</sup>) | 8.4 | 61.7 | 53.8 |
| Year of impoundment | 1980 | 1979 | 1982 |
| Filling time (years) | 0.5 | 1 | 2 |
| Water Residence Time (days) | 124 | 183 | 803 |
| Mean depth (m) | 8 | 22 | 12 |
| Max. depth (m) | 51 | 137 | 49 |
| Mean annual drawdown (m) | 3.6 | 3.3 | 2.1 |
| Max. water temperature (°C) | 20.0 | 20.8 | 20.5 |
| Watershed area (km<sup>2</sup>) | 30.0 | 97.6 | 36.8 |

The waters of the LG complex are naturally characterized by high transparency (1.5 to 4.0 m), high oxygen levels (80% to 100% saturation), slight acidity (pH 5.9–6.9), and low mineral content (conductivity 8–30 µS cm⁻¹). The reservoirs are relatively rich in organic matter but have low nutrient concentrations, with total phosphorus concentrations ranging from 4 to 10 μg L⁻¹. The three reservoirs show marked differences in size and water retention time. For each reservoir, water quality measurements associated with the TSH were collected annually for 15 years following impoundment (Fig. 2).

**Figure 2.**
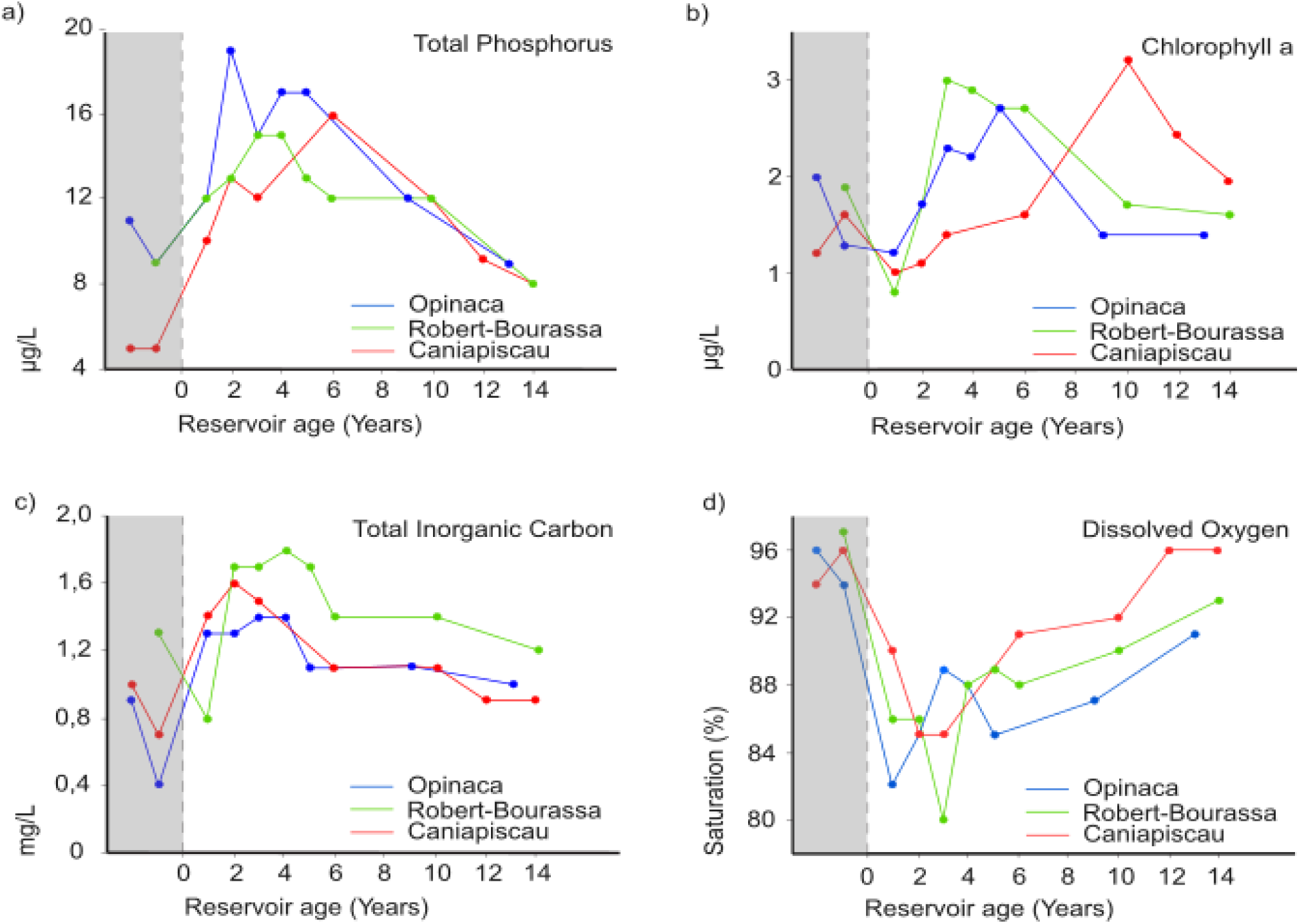
Trends in chlorophyll-a, total phosphorus, total inorganic carbon, and dissolved oxygen saturation in the photic zone during the ice-free period, shown as a function of reservoir age for the three study reservoirs: Robert-Bourassa (red), Opinaca (blue), and Caniapiscau (green). The shaded rectangle in each graph represents the pre-impoundment period. Data extracted from Therrien et al. (2002).

### Data collection

We used Hydro-Québec’s long-term fish community and biogeochemical databases, which stem from annual surveys initiated in 1978 and conducted in July and August, when the reservoirs were free of ice. In the RB reservoir, annual sampling occurred from 1978 to 1984, with additional surveys in 1988, 1992, 1994 (only water quality data), 1996, and 2000. The pre-impoundment period for RB is represented by data collected in 1978. In the OP reservoir annual surveys conducted from 1978 to 1984 were followed by additional sampling in 1988, 1992, 1996, and 2000. The pre-impoundment period for OP spans two years, 1978 and 1979. In the CA reservoir, fish were sampled annually from 1980 to 1982, followed by additional surveys in 1987, 1991, 1993, 1997, and 1999. Pre-impoundment data for CA were collected in 1980 and 1981.

Sampling was conducted at multiple sampling stations in each reservoir: RB: 5 stations, OP and CA: 4 stations. To maintain consistency, station locations were slightly adjusted before and after impoundment to ensure comparable depths, but all stations remained within 500 m of their original positions.

Fish sampling was conducted using two pairs of gill nets set in the littoral zone at depths ranging from 2 to 10 meters. Each pair consisted of one standard experimental net and one uniform-mesh net. The standard nets were 45.7 m long and 2.4 m high, with mesh sizes varying from 2.5 cm to 10.2 cm, whereas the uniform-mesh nets had the same dimensions but had a single mesh size of either 7.6 cm or 10.2 cm. The two pairs of nets were deployed perpendicular to the shoreline in a specific arrangement. For the first pair, the 7.6 cm uniform-mesh net was set closest to the shore with the standard experimental net attached to its offshore end. This experimental net was set with its smallest-mesh section (2.5 cm) nearest the uniform-mesh net and its largest-mesh section (10.2 cm) extending toward the pelagic zone. For the second pair, the standard experimental net was set closest to the shore with its smallest mesh section (2.5 cm) closest to the shore, and the 10.2 cm uniform-mesh net was attached to its offshore end, extending toward the pelagic zone (Therrien et al., 2002). Fish biomass per unit of effort (BPUE) was calculated as kg per net per 24-h period.

Up to 1982, nets were left in place for 48 hours, with retrievals every 24 hours. From 1983 onward, the total soak time was reduced to 24 hours. All captured fish were identified to the species level, measured for total length, and weighed. Details on fish catch data are presented in Table 2.

**Table 2.** Fish species collected in the Robert-Bourassa, Opinaca, and Caniapiscau reservoirs (La Grande complex, northern Quebec). Trophic level is from FishBase (fishbase.se). Species are categorized as either predators (P) or consumers (C) to calculate the predator to consumer mass ratio (PCMR).

| Species (CODE) | Common name | Trophic level | Category |
| --- | --- | --- | --- |
| *Catostomus catostomus* (CACA) | Longnose sucker | 2.5 ± 0.3 | C |
| *Catostomus commersonii* (CACO) | White sucker | 2.8 ± 0.2 | C |
| *Coregonus clupeaformis* (COCL) | Whitefish | 3.2 ± 0.2 | C |
| *Acipenser fulvescens* (ACFU) | Lake sturgeon | 3.3 ± 0.5 | C |
| *Prosopium cylindraceum* (PRCY) | Round whitefish | 3.3 ± 0.4 | C |
| *Salvelinus fontinalis* (SAFO) | Brook trout | 3.3 ± 0.0 | C |
| *Coregonus artedii* (COAR) | Cisco | 3.4 ± 0.4 | C |
| *Perca flavescens* (PEFL) | Yellow perch | 3.7 ± 0.2 | P |
| *Lota lota* (LOLO) | Burbot | 3.8 ± 0.2 | P |
| *Semotilus atromaculatus* (SEMA) | Creek chub | 4.0 ± 0.5 | P |
| *Esox lucius* (ESLU) | Northern pike | 4.1 ± 0.4 | P |
| *Salvelinus namaycush* (SANA) | Lake trout | 4.3 ± 0.5 | P |
| *Sander vitreus* (SAVI) | Walleye | 4.5 ± 0.0 | P |

**Table 3.** Summary output of generalized additive mixed model (GAMM) of the predator–consumer mass ratio (PCMR) as a function of reservoir age for the three study reservoirs (Opinaca, Robert-Bourassa, Caniapiscau).

| Reservoir | Intercept (SE) | Std. error | Effective Degrees of<br>Freedom | F | Adj. R<sup>2</sup> | p value |
| --- | --- | --- | --- | --- | --- | --- |
| Opinaca | 1.334±0.182 | 0.182 | 3.55 | 3.67 | 0.173 | 0.027 |
| Robert-Bourassa | 1.154±0.112 | 0.112 | 4.85 | 10.72 | 0.418 | < 0.001 |
| Caniapiscau | 1.010±0.231 | 0.231 | 1.96 | 5.42 | 0.096 | 0.008 |

### Statistical analyses

We used the PCMR as a metric to capture bioenergetic relationships within fish assemblages (Barnes et al., 2010). By tracking temporal trends in PCMR, we can identify dominant energy pathways and detect shifts in biomass ratios between top predators (piscivores) and other consumer fish. For each reservoir, we calculated the mean PCMR for each sampling station and year by dividing the total predator biomass by the total consumer biomass at each station and year. One outlier from the Opinaca reservoir (PCMR = 107) was excluded from the analyses because it resulted from a low number of prey captures at a single station during the eighth year of monitoring. The classification of species as predators or consumers was based on their trophic position (TP) and dietary habits, using information extracted from FishBase (Froese & Pauly, 2024).

When 0 < PCMR < 1, consumer biomass exceeds predator biomass, reflecting a conventional bottom-heavy trophic pyramid. In contrast, PCMR > 1 signifies that predator biomass surpasses consumer biomass, reflecting an inverted trophic pyramid (McCauley et al., 2018; de Omena et al., 2019). Higher relative PCMR values suggest a bottom-up energy flow, in which the system is primarily driven by energy transfer from lower to higher trophic levels.

Conversely, a reduction in the PCMR ratio reflects top-down regulation, implying that predator dynamics exert primary control over consumer populations (McQueen et al., 1989).

We fitted separate generalized additive mixed models (GAMMs) for each reservoir to examine and compare temporal trends in PCMR across reservoirs. Models were fit using the *gamm* function within the *mgcv* package (version 1.9-4) in R (version 4.5.2). For each reservoir-specific model, the temporal trend in PCMR was modelled as a smooth function of reservoir age (time since impoundment). We used square-root-transformed PCMR (sPCMR) as the dependent variable in the models to improve the normality of the residuals. A gaussian process (bs = “gp”) was used as the smooth function. The basis dimension (*k*) was set to 10 for the OP and RB reservoir models, and to 7 for the CA reservoir model, which required lower dimensionality because it had fewer unique covariate combinations in its dataset. Nesting, and ensuing within-groups correlations arising from repeated sampling at fixed locations, was addressed by including the station indicator as a random effect. Models were fitted using restricted maximum likelihood (REML), which is the preferred method for estimating variance components of the smoothers, as it yields more reliable estimates of the smoothing parameters and balances model fit and smoothness (Wood, 2004, 2017). The R specification for an individual reservoir’s model was: gamm(sPCMR ∼ s(time, bs = “gp”, k = [7 or 10]), random = list(STATION = ∼1), data = [individual_reservoir_data], method = “REML”). After fitting the individual models, the estimated smooth trends were plotted and visually compared to assess differences in the temporal patterns of PCMR among the reservoirs. We used the diagnostics in the *gam.check* function to inspect the adequacy of model fits.

## RESULTS

### Evidence of a trophic surge in La Grande complex reservoirs

The data revealed distinct transient dynamics for total phosphorus (TP), chlorophyll *a*, and total inorganic carbon (TIC), characterized by initial post-impoundment increases followed by decreases to baseline levels (Fig. 2). TP exhibited a consistent unimodal pattern across all three reservoirs, though the timing of the maxima varied. The most rapid and intense peak occurred in OP during year 2, while RB reached its peak in year 3 at a lower concentration than OP. CA displayed the most delayed response, with a peak observed in year 6 (Fig. 2a). Chlorophyll *a* trends largely mirrored those of nutrient dynamics. Concentrations in OP and RB increased within the first four years of impoundment, whereas CA followed a protracted trajectory, reaching its peak approximately 10 years post-impoundment (Fig. 2b). TIC similarly followed a hump-shaped trend, though its decline was more abrupt, leading to stabilization by the year 6 across all systems. RB reached its highest TIC concentration in year 4, while CA peaked earlier in year 2. Following these peaks, TIC levels in all three reservoirs declined and eventually approached pre-impoundment values (Fig. 2c). In contrast, dissolved oxygen (DO) saturation declined sharply and rapidly immediately following impoundment, then recovered steadily. The most pronounced depletion was observed in RB, with OP exhibiting a similar decline. CA showed a less marked decrease and achieved a full recovery to pre-impoundment oxygen levels by year 12 (Fig. 2d).

### Prevalence and magnitude of top-heavy trophic pyramid in reservoirs

In OP and RB reservoirs, PCMR values exceeded 1 quickly after impoundment, indicating ITP. The temporal trends in PCMR showed two distinct patterns across the reservoirs (Fig. 3). OP and RB both followed a clear hump-shaped trajectory, with PCMR increasing to a peak before declining, although the magnitude and duration varied between them. Conversely, the CA reservoir did not follow this pattern. Instead, we observed a steady increase in PCMR throughout the study, with limited evidence of ITP prevalence in CA (values exceeding 1; Fig. 3).

**Figure 3.**
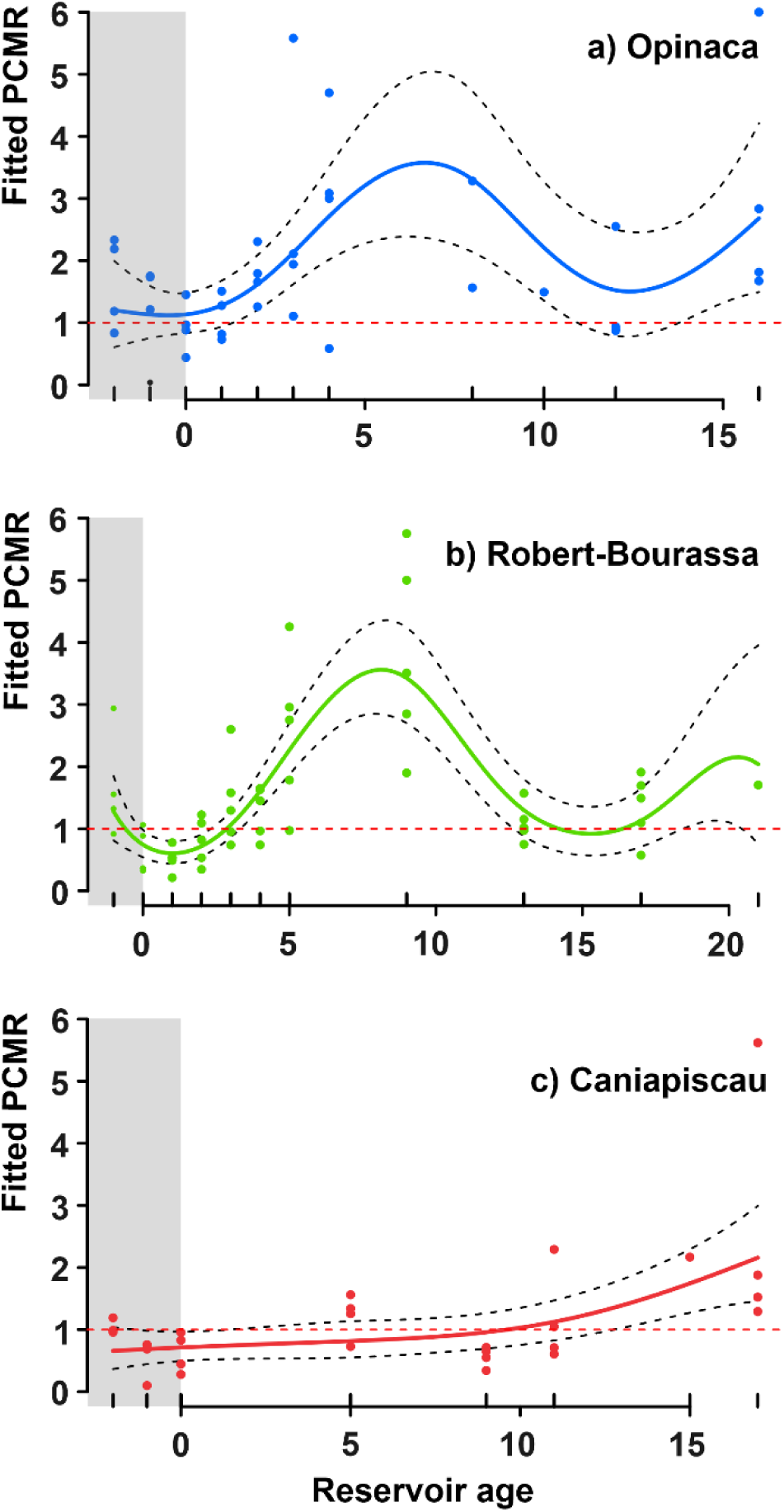
Smoothed trends of the predator-consumer mass ratio (PCMR) over time in the a) Opinaca, b) Robert Bourassa, and c) Caniapiscau reservoirs, located in northern Quebec. Trends were derived using generalized additive mixed models (GAMMs), incorporating time as a smooth term and accounting for variability among sampling stations as random effects. Confidence intervals around the fitted smooths represent 95% confidence intervals. Year 0 denotes the transition point between the pre- and post-impoundment periods. Dashed horizontal line: threshold value associated with the onset of an inverted trophic pyramid.

In the OP reservoir, PCMR values peaked at approximately 3.5 in year 7 and then declined to pre-impoundment levels by year 12. The RB reservoir followed a similar trajectory, with a steeper rise in PCMR values. PCMR values exceeded 1 at year 3, peaked at approximately 3.5 in year 8, and then declined to pre-impoundment levels by year 14. PCMR values were > 1 for about 11 years (from year 3 to 14 post-impoundment; Fig. 3b). The PCMR hump-shaped in OP lasted for about 11 years (from year 1 to 12 post-impoundment; Fig. 3a). In contrast, the CA reservoir exhibited a monotonic upward trend in PCMR values post-impoundment, which accelerated ∼10 years after impoundment and attained mean values greater than 1 for about 7 years (from year 10 to 17 post-impoundment; Fig. 3c).

Diagnostic checks for all reservoirs confirmed that the chosen basis dimension captured the biological trends without evidence of under-fitting (k-index values > 1.15, p > 0.80).

Residual analysis indicated no substantial departures from the assumptions of normality and homoscedasticity.

### Biomass variation of fish species in reservoirs

The restructuring of fish assemblages following impoundment was characterized by shifts in relative biomass for some species across the three reservoirs (Fig. 4). In the OP reservoir, *Esox lucius* (Northern pike) showed a marked increase in dominance, with its relative biomass rising from approximately 10% before impoundment to nearly 60% by year 7. This increase coincided with a decline in consumers of the genus *Catostomus* and a decrease in biomass for *Sander vitreus* (walleye), another top predator. However, walleye, unlike the declining consumer species, began a gradual recovery by year 7 (Fig. 4a). A similar pattern was observed in the RB reservoir, where pike relative biomass increased rapidly following impoundment, rising from 10% before impoundment to nearly 50% within the first 8 years. The decline in *Catostomus* consumers mirrored the decline observed in the OP reservoir. *Coregonus artedi* (cisco) also decreased in relative biomass, contrasting with a concurrent rise in *Coregonus clupeaformis* (whitefish). Walleye exhibited a steady decline throughout the sampling period (Fig. 4b). In the CA reservoir, the patterns of biomass variation resembled those in the OP and RB reservoirs but were more temporally stable. Although pike relative abundance declined in year 9, it resumed an upward trend two years later. As in the OP and RB reservoirs, consumer species — particularly those of the genus *Catostomus* — declined steadily. *Salvelinus namaycush* (Lake trout) maintained a relatively stable relative biomass throughout the study period but increased from 20% to 50% relative to the year of impoundment in the final sampling year (Fig. 4c).

**Figure 4.**
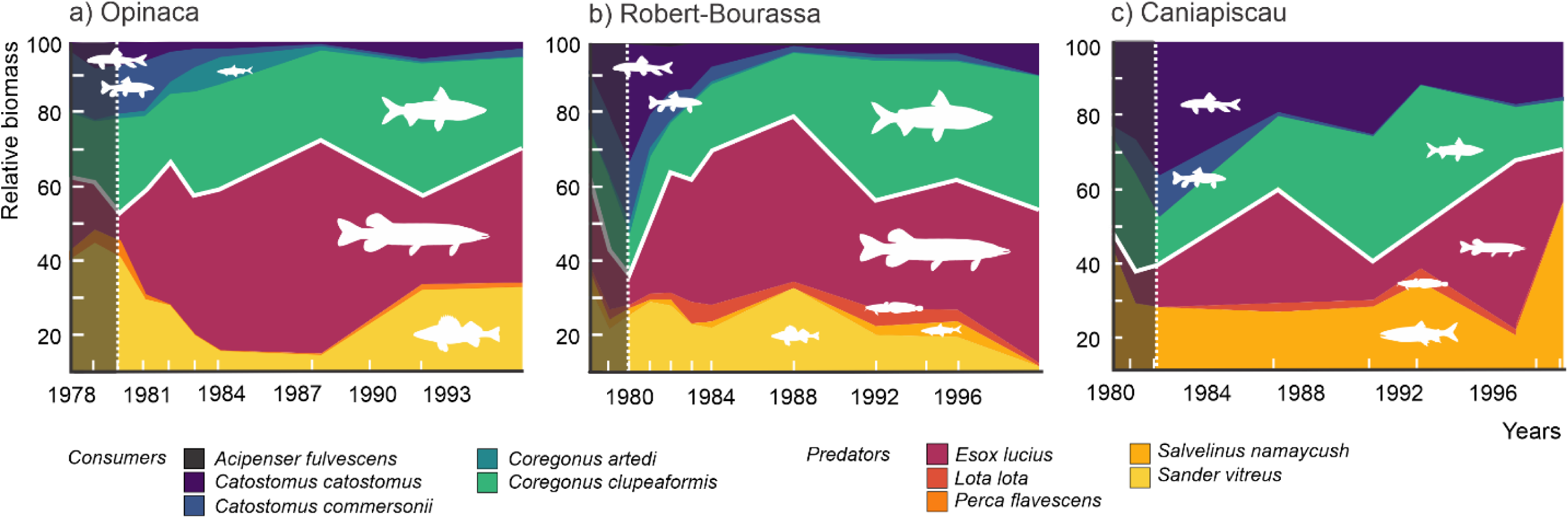
Variation in the relative total biomass of the sampled species over time in the a) Opinaca, b) Robert Bourassa, and c) Caniapiscau reservoirs. Shaded rectangle: pre-impoundment period. The white line separates predators from consumers. The legend displays the color code associated with each species.

## DISCUSSION

Our study shows that the impoundment of large boreal reservoirs leads to a transient reorganization of fish community structure, characterized by the emergence of inverted trophic pyramids (ITP). This top-heavy configuration, where the biomass of predator fish (piscivores) exceeds the biomass of consumer fish (non-piscivorous), seems to be primarily driven by the temporary pulse of nutrients (trophic surge) that sustains productivity across trophic levels.

Trophic disequilibrium, an outcome of the trophic surge, allows populations of large generalist predators, such as *Esox lucius*, to grow faster than the system can sustainably support. Below, we propose a framework that explains how the magnitude and duration of ITPs can be associated with reservoir morphometry and WRT. We suggest that larger reservoirs with longer WRT should exhibit a more gradual and damped transition toward ITP than smaller, faster-flushing reservoirs.

### Trophic Surge: The main driver of top-heavy pyramids in reservoirs

The transient inversion of the fish trophic pyramid in the study reservoirs may result from an imbalance in biomass ratios, triggered by asynchronous and lagged responses of predators and consumers to an intensified bottom-up energy flow during the trophic surge. Given that the OP, RB, and CA reservoirs are naturally oligotrophic, this nutrient influx following impoundment, coupled with prolonged WRT in reservoirs by the dam, boosted primary production, thereby stimulating a bottom-up energy flux during those years (Ostrofsky & Duthie, 1980; Trottier et al., 2024).

TP, chlorophyll *a*, and inorganic carbon displayed a distinct pattern indicative of a trophic upsurge, characterized by a pronounced hump-shaped trajectory (Fig. 2). The delayed peak in chlorophyll *a*, relative to TP, can be attributed to the time required for nutrients to be released from inundated soils and to high turbidity. In newly impounded systems with altered sediment fluxes, turbidity limits light availability and consequently dampens autotrophic uptake during the first few weeks after impoundment (Karlsson et al., 2009; Maavara et al., 2017; Tanabe et al., 2019). The observed decline in oxygen saturation indicates a high rate of organic matter decomposition, which can amplify the mismatch between chlorophyll *a* and TP (Fig. 2d). Even though nutrient release from decomposition fuels phytoplankton blooms, the resulting hypoxia can create a stressful environment and inhibit the growth and development of oxygen-sensitive taxa, such as certain fish and benthic macroinvertebrates (Abbott et al., 2022).

The trophic surge fundamentally reorganizes the food web by boosting the highly efficient, nutrient-rich algae pathway. This surge stimulates secondary productivity—the growth of organisms that consume algae (Cross et al., 2006; Bumpers et al., 2017). As a result, the entire trophic chain benefits from this enhanced food source; for fish in particular, leading to significant changes in their specific diet, biomass distribution, species composition and population densities (Delariva et al., 2013; Pereira et al., 2016, 2021; Turgeon et al., 2016, 2019a; Monaghan et al., 2020; Perônico et al., 2020; Pennock & Gido, 2021). This shift alters the ecosystem’s overall architecture by increasing key biomass ratios (*e.g.*, predator-to-prey) and modifying food-web functions, such as nutrient cycling, regulation of populations, and ecosystem stability (Cebrian et al., 2009; Bumpers et al., 2017; Roussel et al., 2023).

### Transient nature of ITPs in reservoirs and mechanisms of formation

Extending the Trophic Surge Hypothesis (TSH) to top predators helps describe post-impoundment community shifts, though these dynamics are governed by a complex interplay of factors beyond nutrient enrichment alone. Numerous exceptions to the conventional bottom-heavy trophic pyramids have been described in the literature. ITPs emerge as prevailing configurations in ecosystems with energetic pathways influenced by spatio-temporal features that route the flow of matter and energy efficiently (McCauley et al., 2018; Sandin & Zgliczynski, 2015). Although top-heaviness has been reported across ecosystems, it is more frequently observed in aquatic communities (Reuman et al., 2008; McCauley et al., 2018). Biomass ratios exceeding one have been documented in several aquatic ecosystems, at basal trophic levels such as phytoplankton, zooplankton and periphyton communities (Del Giorgio & Gasol, 1995; Vadeboncoeur & Power, 2017), as well as in predator-prey assemblages in reef and oceanic environments (Friedlander & DeMartini, 2002; Sandin & Zgliczynski, 2015; Mourier et al., 2016; Trebilco et al., 2016).

In contrast to such established natural ecosystems which exhibit stable trophic dynamics, top-heaviness in reservoirs seems to be a transient state, triggered as responses to a specific ecological disturbance, namely a nutrient pulse during flooding. In this context, temporal trends in ITPs can serve as indicators of trophic instability, providing a framework to assess the extent and magnitude of damming impacts. We observed a hump-shaped trajectory of PCMR reflecting a temporary increase in top-heaviness in the OP and RB reservoirs. Peak values exceeded 1 and subsequently returned toward their initial values, indicating a transient inversion in the OP and RB reservoirs (Fig. 3). The hump-shaped pattern was not observed in the CA reservoir; however, the increasing trend in PCMR in this reservoir indicates that the community is transitioning toward a top-heavier configuration. Although in the CA reservoir 20 years may not have been sufficient to capture the full transient pattern seen in the OP and RB reservoirs, the contrast in size and water residence time between the OP and RB and the CP reservoirs suggests that the processes driving top-heavy configurations, including trophic pyramid inversions, are influenced by the systems’ capacity to accumulate and retain nutrients from the flooded watershed and upstream waters.

The trophic surge, beyond reshaping energy dynamics within reservoirs, acts alongside other processes that further sustain the transient top-heaviness observed in reservoirs. Overall, ITPs emerge through two main pathways: (1) endogenous mechanisms that enhance energy transfer efficiency or minimize energy losses through digestion, assimilation, and respiration, and (2) exogenous pathways that introduce external energy sources into the community (McCauley et al., 2018). In reservoirs, the trophic surge operates through both mechanisms— initially as an exogenous pathway, with phosphorus input acting as an allochthonous subsidy, and subsequently as an endogenous driver by enhancing primary productivity (Grimard & Jones, 1982). After impoundment, when the ecosystem becomes regulated by the algae-based pathways, primary producers increase both in abundance and nutritional quality, which supports a sustained biomass accumulation across different consumer levels (Ostrofsky & Duthie, 2011). However, restructuring toward a top-heavier configuration requires a prolonged period of high primary productivity with rapid turnover rates (Vadeboncoeur & Power, 2017). In newly impounded reservoirs nutrient availability is high, the continuous renewal of phytoplankton sustains a stable food supply for grazers, and therefore the flux of energy is expected to propagate across successive trophic levels (Heathcote et al., 2016; Vadeboncoeur & Power, 2017). These changes have been linked to higher peaks in fish recruitment, fostering larger adult fish populations and potentially leading to the inversion of the trophic pyramid (Kimmel & Groeger, 1983; Rydin et al., 2008; Parisek et al., 2024).

Our data show that a transient, hump-shaped peak in phosphorus and chlorophyll *a* precedes subsequent shifts in the fish assemblages, as measured by the PCMR, within the three reservoirs. The ITP configurations in reservoirs are temporary, as the underlying resource base is in flux. The system undergoes marked fluctuations between resource surplus to scarcity, leading to a complete reorganization of the food web and its energy-transfer efficiency. Based on this, we propose a conceptual framework (Fig. 5) to explain this predictable transition from an unstable young reservoir to a more mature ecosystem.

**Figure 5.**
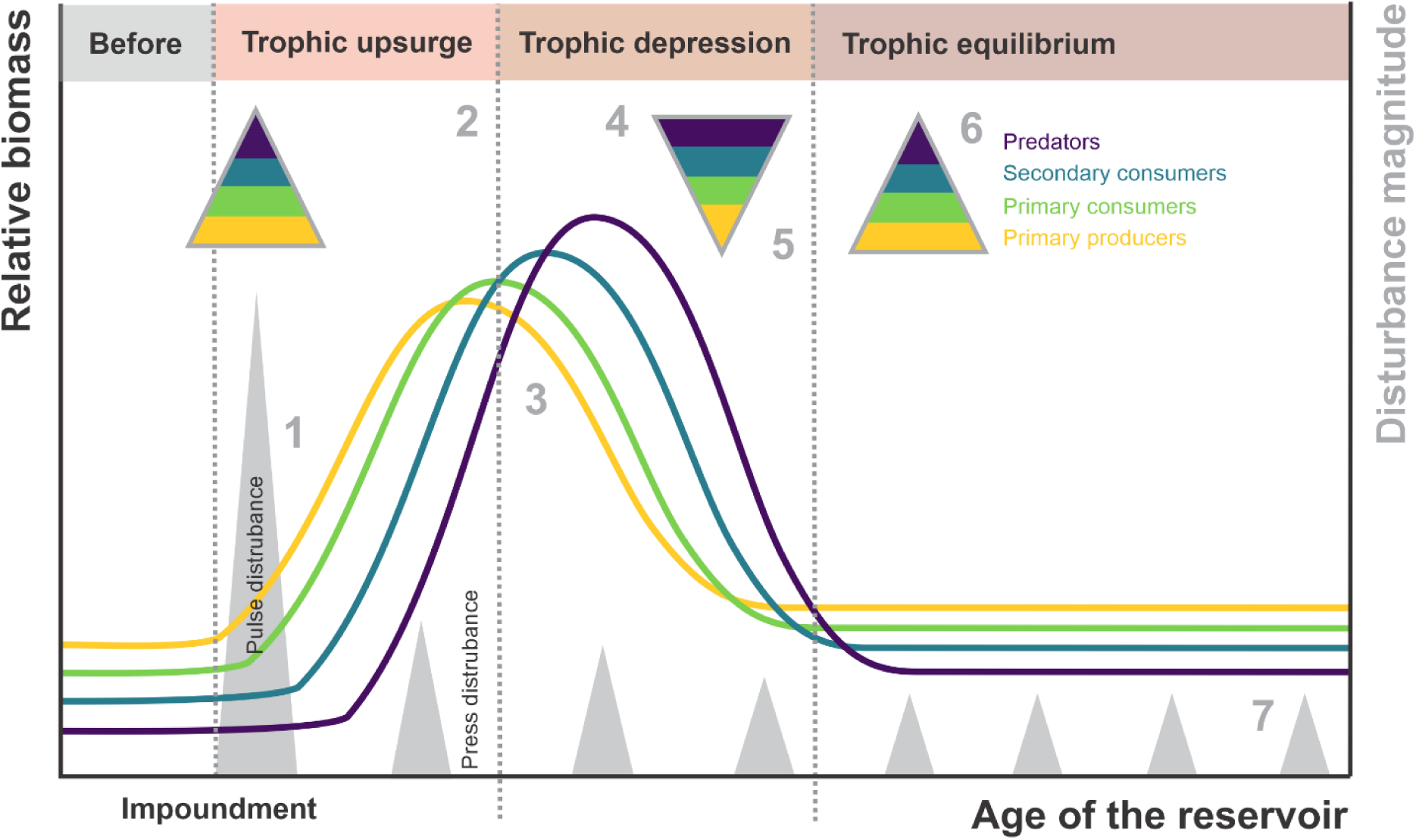
Theoretical framework for the dynamics of inverted trophic pyramids (ITPs) following dam construction. 1. Dam construction, a strong pulse disturbance, drastically alters the hydrology and geomorphology of aquatic ecosystems. 2. Reservoir filling releases nutrients and organic carbon, stimulating primary production and driving a bottom-up trophic regulation that amplifies biomass across trophic levels. 3. As nutrients deplete and inputs decline, primary production begins to decrease. 4. The initial trophic surge disrupts biomass balances, temporarily inverting the trophic pyramid and shifting from bottom-up to top-down regulation. 5. Overpopulation of apex predators causes resource scarcity, intensifying competition, altering predator-prey dynamics, and reorganizing the food web. 6. The system gradually stabilizes, reaching a new dynamic equilibrium in biomass and species composition, influenced by increased ecosystem volume, surface area, and water residence time. 7. Long-term stability depends on structural changes maintained by periodic press disturbances from reservoir-level fluctuations caused by dam operations and management practices.

In agreement with theoretical bioenergetic models (Gellner et al., 2016; Rip & McCann, 2011), our results show that a phase of high productivity leads to stable consumer biomass accumulation and an increase in top-heaviness. The stability declines as predator growth leads to excessive energy and food demands that exceed the system’s productive capacity (Gilbert et al., 2014). In contrast, when resource biomass accumulation is low or consumer growth rates are more moderate, consumer biomass buildup is constrained, promoting stability in trophic interactions—a condition that may resemble the ecosystem’s state prior to dam construction (Gilbert et al., 2014; McCauley et al., 2018). The GAMMs of the PCMR in the OP and RB reservoirs capture both the stabilization and destabilization phases. However, the top-down pressures emerging during destabilization warrant further investigation. Specifically, it remains unclear whether reservoirs attain a dynamic equilibrium once the effects of the trophic surge are regulated or if subsequent press disturbances during dam operation (*e.g.,* fluctuations between flooding and drawdown) sustain the system in oscillatory declines or cyclic population fluctuations.

### Other factors influencing top-heaviness in reservoirs

Our results suggest that the magnitude and duration of top-heaviness may be correlated with reservoir area and water residence time (WRT). The three reservoirs studied are cascade reservoirs and share similar water quality and biodiversity characteristics, as well as several potentially confounding factors, such as low fishing effort and anthropogenic land use. The OP reservoir, which has the smallest area (1,040 km²) and the shortest residence time (124 days), exhibited a transient top-heavy pulse that peaked between years 5 and 10 with a maximum PCMR of approximately 3.5. Similarly, the RB reservoir, with a larger area (2,835 km²) and longer residence time (183 days), experienced a top-heavy phase that also peaked near a PCMR of 3.5 around year 8 before declining. The CA reservoir, which has the largest area (4,275 km²) and longest residence time (803 days), showed a steady increase in top-heaviness over the 17-year sampling period, with the PCMR consistently rising above baseline, exceeding 2 by the end of the study period.

Geomorphological properties are fundamental drivers of food web architecture and stability in aquatic ecosystems (Cohen & Newman, 1988; Post et al., 2000). Understanding the influence of these properties on the transient trophic dynamics of reservoirs is essential for improving the design of reservoir creation to mitigate impacts on biotic communities. The ecological function, carrying capacity, and biological productivity of reservoirs are largely determined by their post-impoundment morphometry. Morphometry plays a key role in regulating processes such as nutrient cycling, sedimentation, mixing, temperature regimes, and trophic interactions (Fee, 1979; Håkanson, 2005). In the context of trophic productivity, reservoir area is particularly influential, as it dictates the capacity to receive and redistribute nutrients and dissolved organic carbon after flooding and the efficiency of energy transfer between macrohabitats (*i.e.,* pelagic-littoral-benthic; Guildford et al., 1994; McCann et al., 2005). WRT determines the extent to which a reservoir retains or flushes nutrients downstream, thereby influencing nutrient budgets and species succession (Ambrosetti et al., 2003; Rueda et al., 2006; Tong et al., 2019). Therefore, the trophic stability in large reservoirs, such as the CA reservoir, may take decades to reach equilibrium after dam construction and impoundment (Prchalová et al., 2009; Říha et al., 2009).

Changes in physical habitat and hydrological regime induced by impoundment are a primary driver of fish community structure, creating conditions that favor the proliferation of certain species while restricting others (Agostinho et al., 2008; Poff et al., 2007). As the riverine environment transitions to a lake-like environment, evolutionarily established communities are disrupted. Generalist species adapted to slow-moving, open-water (lentic) environments often thrive, while species requiring fast-flowing (lotic) habitats decline (Poff et al., 2007). The establishment of a dynamic equilibrium in the new reservoir largely depends on how different species respond to these new morphometric conditions (Swales, 2006; Poff et al., 2007).

Theoretical trophic network analyses suggest that smaller lakes exhibit greater coupling between pelagic and littoral zones, allowing highly mobile pelagic predators greater access to littoral resources (Dolson et al., 2009; Tunney et al., 2012). This increased resource accessibility often leads to food chain contraction and a top-heavy biomass pyramid (McCann et al., 2005).

Additionally, WRT modulates top-heaviness by influencing physical processes, such as the transport and mixing of dissolved and suspended particles and nutrients, and more generally regulates trophic state, thermal stratification, isotopic composition, alkalinity, heavy metal and nutrient ratios, organic matter mineralization rates, primary production, and predator-prey interactions (Ambrosetti et al., 2003; Rueda et al., 2006). Reservoirs with longer residence times are more susceptible to stronger and more frequent algal blooms (Gibson et al., 2000). As reservoirs trap resources from their watersheds, longer WRT allow these resources to be processed into bioavailable forms throughout the trophic chain, increasing transfer efficiency and promoting top-heaviness (Burford et al., 2007).

In recently impounded reservoirs, the physical habitat is reconfigured and lacks the stable, compartmentalized zones of mature lakes (e.g., pelagic, littoral, and benthic zones; Drakou et al., 2008). The initial flooding of the landscape creates a complex mosaic of new niches, particularly along the reservoir edges, which alters predator-prey dynamics by providing new refuges for prey (Wang et al., 2009). Fish communities undergo restructuring in response to a suite of environmental changes, from water temperature and oxygen levels to prey availability and competition (Gido et al., 2000; Brosse et al., 2007; Říha et al., 2009; Pennock & Gido, 2021). The resulting community is shaped by species-specific adaptations—balancing the need to acquire resources with the risk of predation in the new and evolving environment (Prchalová et al., 2009).

The hump-shaped trajectory of the PCMR values in the La Grande reservoirs closely mirrored the trends in pike (*Esox lucius*) relative biomass (Fig. 4). As a top predator, pike plays a crucial role in structuring fish communities (Craig, 2008). Reservoirs provide favorable environmental conditions for pike (*i.e.*, calm waters and abundant flooded vegetation), which increase hunting success and the availability of spawning habitats (Forsman et al., 2015). *E. lucius* is a generalist species that quickly adapts to reservoirs and can undergo ontogenetic niche shifts, particularly in diet, which results in resource partitioning when competing with other predators (Amundsen et al., 2003; Westrelin et al., 2022), or cannibalism, allowing high biomass accumulation in large individuals (Craig, 2008). The capacity of *E. lucius* to exploit the resources available in reservoirs might make this species a driver of top-heaviness in the La Grande reservoirs.

### Methodological considerations and constraints

While the longitudinal consistency of the Hydro-Québec monitoring program provides an invaluable baseline for long-term studies in reservoirs, several ecological and technical constraints warrant consideration. First, the reliance on experimental gill nets can potentially introduce size and activity-dependent biases (Hamley, 1975). For instance, gillnets can be selective for shape and size of predators such as *Esox lucius*, particularly when exploiting new hunting grounds and spawning habitats in recently flooded littoral zones, which may lead to their over-representation in catches compared to more sedentary benthic consumers like *Catostomus* species. Although the use of standardized multifilament nets across all years ensures internal consistency, the magnitude of the observed shifts in the PCMR suggests a fundamental community reorganization that transcends sampling artifacts.

Furthermore, the calculation of the PCMR utilized binary “predator” or “consumer” assignments based on established adult trophic positions (TP). This approach provides a clear metric for biomass distribution but simplifies complex ontogenetic niche shifts in which species like *Perca flavescens* or *Sander vitreus* transition from primary consumers to apex predators as they increase in size.

## CONCLUSIONS

The transient inversion of the trophic pyramid, or top-heaviness, observed in three oligotrophic reservoirs of the La Grande complex may result from a trophic surge driven by nutrient pulses from flooded soils following impoundment. This imbalance in biomass ratios, characterized by a bottom-up surge in primary production, triggers a shift in fish community structure, increasing the biomass of piscivorous predators before transitioning to top-down regulation. Chlorophyll *a*, phosphorus, and inorganic carbon concentrations reveal a clear trophic surge with a hump-shaped trajectory, indicating a peak in primary productivity, followed by a decline and eventual stabilization of the ecosystem. Apex predators, such as *E. lucius*, appeared to play a major role in shaping fish community structure. *E. lucius* capitalizes quickly on the resources and habitat conditions created by impoundment and exerts a strong influence on the development of top-heaviness, reinforcing its role as a primary driver of trophic dynamics in these reservoirs. Reservoir area and WRT might be related to the magnitude, timing, and duration of ITP in our study. This observation underscores the critical role of reservoir morphometry and nutrient retention capacity in shaping trophic dynamics and fish productivity. In summary, the trophic upsurge and resulting top-heaviness observed in the La Grande reservoirs provide a clear example of how dams, by modifying hydrological and nutrient dynamics, reshape species interactions in aquatic food webs. The transient nature of ITPs in these systems underscores the need for long-term monitoring to elucidate the factors governing their stability and productivity.

## Conflict of interest

The authors declare that the research was conducted in the absence of any commercial or financial relationships that could be construed as a potential conflict of interest.

## Data availability statement

The databases used in this study are owned by Hydro-Québec and are not publicly available due to confidentiality restrictions. Data may be available upon request with the explicit permission of Hydro-Québec.

## Author contributions

Conceptualization: AS-C, KT; Data curation: AS-C; Formal analysis: AS-C, KT, MAR; Funding acquisition: KT; Investigation: AS-C; Methodology: AS-C, KT, MAR; Project administration: AS-C, KT; Resources: AS-C, KT; Software: AS-C, MAR; Super-vision: KT, MAR; Validation: AS-C, KT, MAR; Visualization: AS-C, KT, MAR; Writing – original draft: AS-C; Writing – review & editing: AS-C, KT, MAR.

## ACKNOWLEDGEMENTS

This work was supported by funding from Hydro-Québec via a Mitacs Acceleration grant. ASC received scholarship support from the Fonds de recherche du Québec – Nature et technologies (merit scholarship program for foreign students, PBEEE, DOI: https://doi.org/10.69777/332321).

## Notes

### Competing Interest Statement

The authors have declared no competing interest.

